# Prolonged oral coenzyme Q_10_-β-cyclodextrin supplementation increases plasma CoQ_10_ concentration and skeletal muscle complex I+III activity in young, untrained healthy Thoroughbreds

**DOI:** 10.1101/818120

**Authors:** Mary F. Rooney, Caitriona E. Curley, James Sweeney, Michael E. Griffin, Richard K. Porter, Emmeline W. Hill, Lisa M. Katz

## Abstract

Coenzyme Q_10_ (CoQ_10_) is an essential component of the mitochondrial electron transport chain (ETC). Decreased skeletal muscle CoQ_10_ content may result in decreased ETC activity and energy production. This study aimed to test the hypothesis that prolonged supplementation with oral CoQ_10_ will increase plasma CoQ_10_ concentrations and skeletal muscle CoQ_10_ content in young, healthy untrained Thoroughbreds. Nineteen Thoroughbreds (27.5±9.7 months old; 11 males, 8 females) from one farm and maintained on a grass pasture with one grain meal per day were supplemented orally once per day for 9 weeks with 1.5 mg/kg body weight of a CoQ_10_-β-cyclodextrin inclusion complex. Whole-blood and skeletal muscle biopsies were collected before (T_0_) and after (T_1_) 9 weeks of supplementation. Plasma CoQ_10_ concentrations were determined via high-performance liquid chromatography. Skeletal muscle mitochondrial ETC combined complex I+III enzyme activity (an indirect measurement of CoQ_10_ content) was assessed spectrophotometrically and normalised to mitochondrial abundance. Results were analysed using a paired two-tailed Students *t*-test with *P*≤0.05 significant. Horses accepted supplementation with no adverse effects. The mean change in plasma CoQ_10_ concentration from T_0_ to T_1_ was significantly greater than zero (0.13±0.02 *vs*. 0.25±0.03 µg/ml, mean difference 0.12±0.03; *P*=0.004), although variability in absorbance resulted in only a 58% response rate. The mean change in skeletal muscle complex I+III activity from T_0_ to T_1_ was significantly greater than zero (0.36±0.04 *vs*. 0.59±0.05 pmol/min/mg of muscle, mean difference 0.23±0.05; *P*=0.0004), although T_1_ values for 3/19 horses decreased on average by 23% below T_0_ values. In conclusion, prolonged oral supplementation of the diet of young, healthy untrained Thoroughbreds with CoQ_10_ increased mean plasma CoQ_10_ concentration by 99% and mean skeletal muscle complex I+III activity by 65% with variability in absorbance among horses. Additional research is warranted investigating training and exercise effects on skeletal muscle CoQ_10_ content in CoQ_10_ supplemented and un-supplemented Thoroughbreds.

## Introduction

Coenzyme Q_10_ (CoQ_10_, ubiquinone) is a small lipophilic molecule endogenously synthesised (Olson & Rudney, 1983; Tran & Clarke, 2007) in eukaryotic cells with a principal role in aerobic respiration. CoQ_10_ is a mobile component of the electron transport chain (ETC) within the mitochondrial inner membrane where it transfers electrons from NADH:ubiquinone oxidoreductase (complex I) and succinate dehydrogenase (complex II) to ubiquinol-cytochrome *c* oxidoreductase (complex III) (Mas & Mori, 2010).

In humans, chronic diseases such as chronic heart failure, hypertension and Parkinson’s disease are characterised by low plasma concentration and tissue CoQ_10_ content, with CoQ_10_ supplementation shown to improve clinical responses to treatment (Hofman-Bang, Rehnqvist, Swedberg, Wiklund, & Åström, 1995; Jankowski, Korzeniowska, Cieślewicz, & Jabłecka, 2016; Mortensen et al., 2014; Yang et al., 2015). Healthy human athletes have also been found to develop CoQ_10_ deficiencies, believed to be due to increased metabolic demand (Cooke et al., 2008; M. Kon et al., 2007; Orlando et al., 2018; Zhou, Zhang, Davie, & Marshall-Gradisnik, 2005). Deficiencies in skeletal muscle CoQ_10_ are thought to result in less efficient energy transduction due to decreased ETC activity and suboptimal ATP production (Lenaz et al., 1999), resulting in reduced effective skeletal muscle contractile function and earlier onset of fatigue (Cooke et al., 2008; M. Kon et al., 2007; Michihiro Kon et al., 2008; Kwong et al., 2002; Mizuno et al., 2008). Numerous studies support CoQ_10_ supplementation in human athletes to improve exercise capacity, aerobic power and recovery after exercise (Alf, Schmidt, & Siebrecht, 2013; Bonetti, Solito, Carmosino, Bargossi, & Fiorella, 2000; Cooke et al., 2008; Leelarungrayub, Sawattikanon, Klaphajone, Pothongsunan, & Bloomer, 2010; Mizuno et al., 2008).

Approximately 50% of the total body CoQ_10_ in humans is found in the mitochondrial inner membrane (Greenberg & Frishman, 1990; Kumar, Kaur, Devi, & Mohan, 2009), with organs containing large numbers of mitochondria such as skeletal muscle having the largest amount of CoQ_10_. In healthy people, plasma CoQ_10_ concentrations are always higher than skeletal muscle with any movement of CoQ_10_ from plasma into skeletal muscle due to simple diffusion. The rate of CoQ_10_ movement from the plasma into the mitochondrial inner membrane is limited by the large size and lipophilic nature of this molecule (Kaikkonen, Tuomainen, Nyyssönen, & Salonen, 2002; Turunen, Swiezewska, Chojnacki, Sindelar, & Dallner, 2002), with movement of exogenous CoQ_10_ into most tissues other than plasma previously believed to be low-to-absent (Svensson et al., 1999; Zhou et al., 2005). However, oral CoQ_10_ supplementation has been shown to elevate CoQ_10_ skeletal muscle mitochondrial content in rodents and humans (Cooke et al., 2008; Kamzalov, Sumien, Forster, & Sohal, 2003; M. Kon et al., 2007; Linnane et al., 2002). Although plasma CoQ_10_ concentrations can easily be measured, this only reflects the bioavailability of CoQ_10_ after oral supplementation and not the amount of uptake into skeletal muscle (Duncan et al., 2005; Zhang, Aberg, & Appelkvist, 1995). CoQ_10_ has not been extensively researched in horses, with a few studies demonstrating oral CoQ_10_ supplementation to increase plasma concentrations (Sinatra, Chopra, Jankowitz, Horohov, & Bhagavan, 2013; Sinatra, Jankowitz, Chopra, & Bhagavan, 2013).

The aim of this study was therefore to test the hypothesis that prolonged oral supplementation of CoQ_10_ to the established diet of a group of young, healthy untrained Thoroughbreds would increase plasma CoQ_10_ concentrations and skeletal muscle CoQ_10_ content.

## Materials and Methods

### Animals and experimental design

This study took place from the last week of May ─ end of July 2017 in the Republic of Ireland. Approval was obtained from the University College Dublin Animal Research Ethics Committee with informed owner consent. The project was licenced under the Health Products Regulatory Authority (Ireland).

Nineteen clinically healthy and privately-owned Thoroughbreds from one farm (11 intact males [mean age 27.8±9.0 months], 8 females [mean age 27.1±11.1 months]) that were not currently and had never been in an exercise training programme were included into the study. Prior to and during the entire study, all horses had been/were maintained full-time in small groups of 5─6 horses on 5-acre grass pastures located next to each other. The diet of the horses at the time of entering into the study and during the study consisted of free-choice pasture grazing and one grain meal (one standard scoop of mixed oats) given in the morning. All horses had physical examinations, haematology, biochemistry and faecal evaluations performed prior to inclusion into the study. Body weight (BW) was estimated for each horse at the beginning of the study using a weight tape and formula (Carroll & Huntington, 1988): BW (kg)=[Girth^2^(cm) × Length (cm)]/11,877.

Each horse acted as its own control with jugular whole-blood and skeletal muscle biopsy samples taken before (T_0_) and after 9 weeks (T_1_) of oral CoQ_10_ supplementation to their established diet. All horses were supplemented and sampled during the same 9-week period. A dose of approximately 1.5 mg/kg BW of a CoQ_10_-β-cyclodextrin inclusion complex in powder form (MicroActive® CoQ_10_, Maypro Industries, New York, USA) containing 26% CoQ_10_ w/w was used. For a 500 kg BW horse, this equated to a daily amount of approximately 200 mg of CoQ_10_. The supplement was dissolved in water and administered via syringe immediately after the morning grain between 7–8 am. All blood and skeletal muscle biopsy samples were taken between 11 am–1 pm, 4–6 hrs after the morning grain. T_0_ samples were taken the day before oral CoQ_10_ supplementation had begun with T_1_ samples taken on the last day of oral CoQ_10_ supplementation.

### Sample collection

Jugular venous whole-blood samples were collected from each horse into a lithium heparin vacutainer for measurement of plasma CoQ_10_ concentrations. Plasma was separated from whole-blood within 3 hrs of collection via centrifugation (1,500 g for 5 mins) and stored at −20°C until batch analysis.

Skeletal muscle biopsies were taken from the middle gluteal muscle from standing un-sedated horses as previously described by Ledwith and McGowan (2004). Once collected, all samples were immediately stored on dry ice for transport to the laboratory (within 3 hrs of collection) and subsequently stored at −70°C until analysis.

### Quantification of plasma CoQ_10_ concentrations

Plasma CoQ_10_ concentrations were measured using a validated reverse-phase high-performance liquid chromatography (HPLC) assay by CAL Ltd (Dublin, Ireland) for a randomly chosen sub-set (*n*=12) of the study horses (www.randomizer.org). Plasma samples were extracted by liquid:liquid extraction (ethanol:methanol; 45:55, v/v) on a Synergi C_18_ column with detection carried out at 275 nm with a UV detector. Each plasma sample was assayed in triplicate under oxidized conditions for total CoQ_10_ (ubiquinone+ubiquinol) content. Plasma CoQ_10_ concentrations were calculated from a standard curve produced by standard CoQ_10_ (Sigma-Aldrich, Co. Wicklow, Ireland) in the concentration range 0.156–2.50 µg/ml.

### Quantification of skeletal muscle CoQ_10_ content

Skeletal muscle CoQ_10_ content was measured spectrophotometrically by combined complex I+III assay (indirect measure of CoQ_10_). All reagents were purchased from Sigma-Aldrich (Co. Wicklow, Ireland) unless stated otherwise. Enzyme activity assays were performed at 30°C on a Libra S12 spectrophotometer (Biochrom Ltd., Cambridge, UK) with absorbance changes measured using an attached chart recorder. The activity of each enzyme was measured in triplicate on the same homogenate for each sample.

#### Preparation of skeletal muscle homogenates

Skeletal muscle homogenates were prepared from tissue stored at −70°C. Any fat/connective tissue was removed from the sample before it was weighed using a fine balance (ME104 Mettler Toledo [Mason Technology, Dublin, Ireland], 0.08 mg repeatability). The tissue was then homogenised using an Ultra Turrax T25 (Janke & Kunkel IKA-Labortechnik, Staufen, Germany) in sucrose muscle homogenisation buffer (20mM tris-HCl, 40mM KCl, 2 mM EGTA, 250 mM sucrose, 1 mM ATP, 5 mM MgCl_2_, pH 7.4). An aliquot of the sample was used to perform protein determination using the bicinchoninic acid assay as described by Smith et al. (1985).

#### Citrate synthase activity assay

Citrate synthase enzyme activity (a measure of mitochondrial abundance) was measured spectrophotometrically by a coloured coupled reaction, using a method adapted from Srere (1969). The activity of citrate synthase was determined by monitoring the rate of production of thionitrobenzoic acid at a wavelength of 412 nm. Skeletal muscle homogenate (approximately 5 μg) was incubated in a 1 ml cuvette with tris buffer (0.2 M, pH 8.1) with reaction components 5,5’-dithiobis-(2-nitrobenzoic acid) (0.1 mM), acetyl coenzyme A (0.3 mM) and Triton X (0.1%) added. A blank rate was measured for 2 mins before oxaloacetate (0.5 mM) was added to initiate the reaction with any increase in absorbance monitored for 3 mins. Specific enzyme activity was expressed as pmol/min/mg of muscle protein using the molar extinction coefficient 13,600 L/mol/cm for citrate synthase at 412 nm.

#### NADH cytochrome c oxidoreductase (Complex I+III) activity assay

The activity of NADH cytochrome *c* oxidoreductase (Complex I+III) is an indirect measure of CoQ_10_. As part of the Q cycle in mitochondria, CoQ_10_ transfers electrons from complex I and complex II to complex III. Thus, measurement of combined complex I+III activity gives an indirect measure of CoQ_10_ content, as the activity of these two complexes in combination is dependent on CoQ_10_ (Lerman-Sagie et al., 2001; Leshinsky-Silver et al., 2003). Complex I+III activity was determined in the present study by monitoring the reduction of cytochrome c at 550 nm as per the method described by Powers et al. (2007). Homogenate samples (approximately 20 μg) were incubated in distilled H_2_O in a 1 ml cuvette to allow osmotic shock to occur. After 2 mins incubation, the reaction components potassium phosphate pH 7.5 (50 mM), oxidised cytochrome *c* (50 μM), KCN (0.3 mM), and fatty-acid free BSA (1 mg/ml) were added; a blank rate was measured for 2 mins. NADH (0.2 mM) was then added to initiate the reaction with any increase in absorbance monitored for 3 mins. Following this, rotenone (10 μM) was added and the rate monitored for a further 2 mins. Complex I+III combined specific activity was taken as the rotenone-sensitive activity determined by subtracting the rotenone-resistant activity from the total activity. Specific enzyme activity for complex I+III was expressed as pmol/min/mg of muscle protein using the molar extinction coefficient 18,500 L/mol/cm for reduced cytochrome *c* at 550 nm. Complex I+III activity was subsequently expressed as a ratio to citrate synthase activity to account for the mitochondrial enrichment of the skeletal muscle homogenates.

### Statistical analysis

Statistical analyses were performed using R 3.3.2 (R Foundation for Statistical Computing, Vienna, Austria). The effects of sex and age were investigated using multivariable linear regression models with interaction effects included for age and sex. Baseline age was set at 13 months (the minimum age of horses in the dataset). Horse ages were subsequently not adjusted between measurement time-points as age was not identified as a significant factor in T_0_ plasma values. Where age and sex effects were deemed non-significant and excluded from the model, mean values were compared using a paired two-tailed Students *t*-test with 95% confidence intervals. Spearman’s rank correlation was performed to assess correlation between plasma CoQ_10_ concentrations and skeletal muscle CoQ_10_ content. A *P*≤0.05 indicated significance, with all results expressed as mean ± SEM unless otherwise indicated.

## Results

All horses readily accepted CoQ_10_ supplementation with no adverse effects observed. Descriptive statistics are summarised in Table 1 and 2.

**Table 1.** Summary statistics for plasma CoQ_10_ concentrations measured in triplicate from *n*=12 young, healthy untrained Thoroughbred horses before (T_0_) and after (T_1_) 9 weeks of daily oral supplementation of the established diet with CoQ_10_ (CoQ_10_-β-cyclodextrin complex with 26% CoQ_10_, w/w).

| $n=12$ | T<sub>0</sub><br>($\mu\text{g/ml}$) | T<sub>1</sub><br>($\mu\text{g/ml}$) |
| --- | --- | --- |
| Minimum | 0.04 | 0.12 |
| 25% percentile | 0.08 | 0.14 |
| Median | 0.11 | 0.27 |
| 75% percentile | 0.14 | 0.29 |
| Maximum | 0.33 | 0.49 |
| Mean | 0.13 | 0.25<sup>†</sup> |
| Standard deviation | 0.08 | 0.11 |
| Standard error of the mean | 0.02 | 0.03 |
| 95% Confidence Intervals | 0.08–0.17 | 0.18–0.32 |

<sup>†</sup>denotes significant difference from T<sub>0</sub> values (paired two-tailed Student's $t$ -test, $P \leq 0.01$ ).

**Table 2.** Middle gluteal skeletal muscle CoQ_10_ content for 19 young, healthy untrained Thoroughbred horses before (T_0_) and after (T_1_) 9 weeks of daily oral supplementation of the established diet with CoQ_10_ (CoQ_10_-β-cyclodextrin complex with 26% CoQ_10_, w/w). CoQ_10_ content was assessed by spectrophotometrically measuring skeletal muscle mitochondrial complex I+III activity. Complex I+III activity data (pmol/min/mg of muscle protein) was normalised to mitochondrial abundance (citrate synthase activity)/g of skeletal muscle.

| *n*=19 | T<sub>0</sub><br>(pmol/min/mg muscle protein) | T<sub>1</sub><br>(pmol/min/mg muscle protein) |
| --- | --- | --- |
| Minimum | 0.11 | 0.18 |
| 25% percentile | 0.23 | 0.44 |
| Median | 0.36 | 0.63 |
| 75% percentile | 0.51 | 0.8 |
| Maximum | 0.56 | 0.92 |
| Mean | 0.36 | 0.59<sup>†</sup> |
| Standard deviation | 0.16 | 0.22 |
| Standard error of the mean | 0.04 | 0.05 |
| 95% Confidence Intervals | 0.28–0.43 | 0.49–0.7 |

<sup>†</sup>denotes significant difference from T<sub>0</sub> values (paired two-tailed Student's *t*-test, $P \leq 0.01$ ).

Multivariable linear models were used to evaluate for interactions between age, sex and plasma CoQ_10_ values. Males had higher T_0_ plasma CoQ_10_ values than females (*P*=0.009), with no differences in T_1_ values. For females, age was only significantly associated with T_1_ plasma CoQ_10_ values, with increasing age associated with increasing values (*P*=0.02). For males, increasing age was significantly associated with reductions in T_0_ (*P*=0.03) and T_1_ plasma CoQ_10_ values (*P*=0.02). These results are all tenuous, however, since a single elevated T_0_ plasma CoQ_10_ value for a 13-month-old male horse skewed all statistical outcomes. For the paired differences in plasma CoQ_10_ values between T_0_ and T_1_, a multivariable model including age and sex as factors identified increasing age to be linked to increasing plasma CoQ_10_ values (*P*=0.02). However, the paired differences for plasma CoQ_10_ values between time-points were not significantly associated with sex (males *P*=0.07, females *P*=0.45). When sex was subsequently excluded from the model, the significant association between age and plasma CoQ_10_ values was lost (*P*=0.06). It appears that inadequate power (*n*=12) did not allow completely accurate statistical evaluation of sex and age effects on plasma CoQ_10_ values.

The T_0_ and T_1_ intra-assay coefficient of variations were 13.3% and 5.7%, respectively. The average T_1_ plasma CoQ_10_ concentrations significantly increased by 99% above the average T_0_ measurements (0.13±0.02 µg/ml *vs.* 0.25±0.03 µg/ml, mean difference 0.12±0.03; *P*=0.004; Table 1, Figure 1). Although the T_1_ plasma CoQ_10_ concentrations were higher than the T_0_ measurement for all horses with an average mean of the ratios (i.e., the average of each individual horse’s difference between T_0_ and T_1_ values) showing a 162% of an increase of T_1_ values above T_0_ values, there was a large amount of individual variation ranging from a 0.6– 617.4% of an increase above T_0_ values. Using a measure of uniform bioavailability defined as at least a doubling of T_1_ plasma CoQ_10_ concentrations above T_0_ values, there was a 58% response rate with 7/12 horses meeting this threshold.

**Figure 1.**
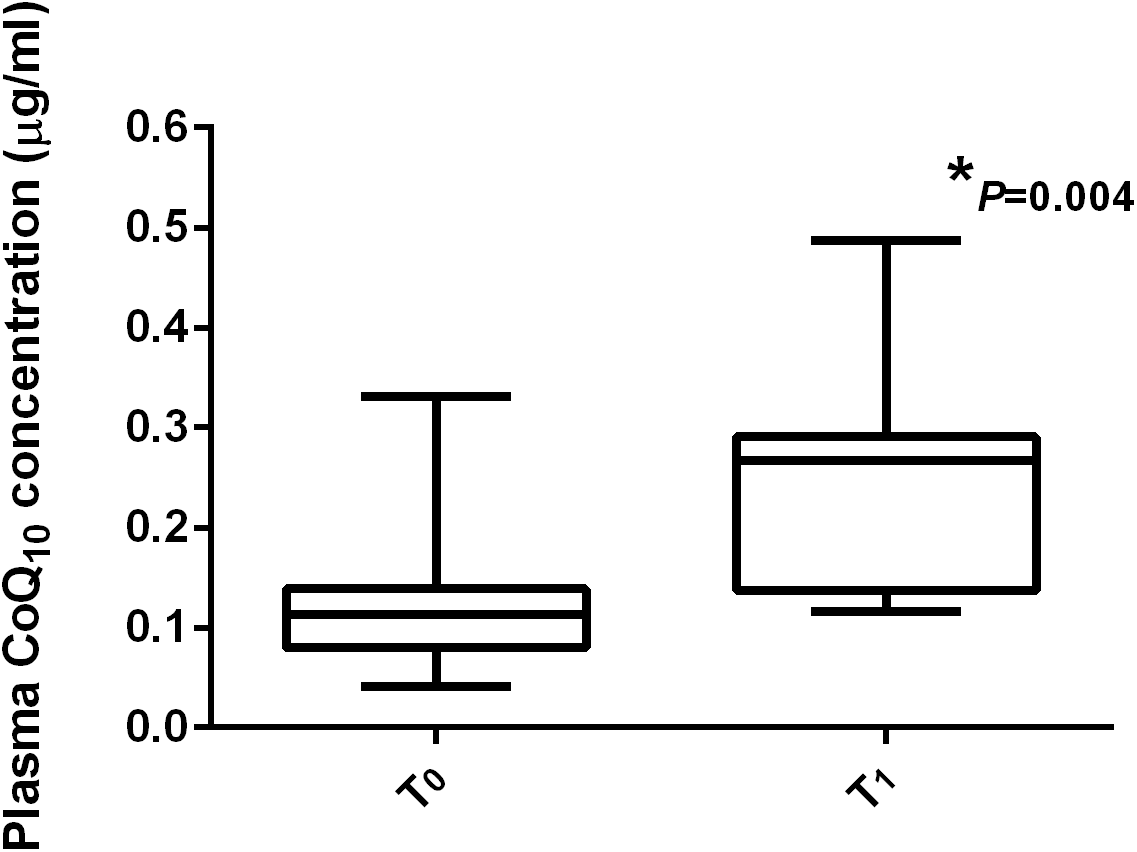
Plasma CoQ_10_ concentrations for 12 young, healthy untrained Thoroughbred horses before (T_0_) and after (T_1_) 9 weeks of daily oral supplementation of the established diet with CoQ_10_ (CoQ_10_-β-cyclodextrin complex with 26% CoQ_10_, w/w). **\***Significantly different from T_0_ values (paired two-tailed Student’s *t*-test, *P*≤0.01).

Multivariable linear models were used to evaluate for interactions between age, sex and skeletal muscle complex I+III activity. Age (*P*=0.84) and sex (*P*=0.06) were not significantly associated with mean T_0_ skeletal muscle complex I+III activity. Age (*P*=0.75) and sex (*P*=0.30) were also not significantly associated with T_1_ skeletal muscle complex I+III activity. For the paired differences in skeletal muscle complex I+III activity between T_0_ and T_1_, neither age (*P*=0.98) nor sex (*P*=0.81) were significantly associated with skeletal muscle complex I+III activity. These results support that any change in mean skeletal muscle complex I+III activity between time-points is independent of both age and sex.

No differences in citrate synthase activity were observed between T_0_ and T_1_ time-points. The average T_1_ skeletal muscle CoQ_10_ content significantly increased above T_0_ values by 65.1% (0.36±0.04 *vs* 0.59±0.05 pmol/min/mg of muscle protein, activity normalised to mitochondrial abundance/g muscle, mean difference 0.23±0.05; *P*=0.0004; Table 2, Figure 2). For 16/19 horses, T_1_ skeletal CoQ_10_ content had increased on average 85% above T_0_ values with a degree of variation ranging from a 13.3–420.9% of an increase above T_0_ values. However, for 3/19 horses, T_1_ skeletal CoQ_10_ content decreased by an average of 22.7% (range 11.4–32.4% of a decrease) below T_0_ values. There were no correlations between T_0_ plasma and skeletal muscle CoQ_10_ measurements nor between T_1_ plasma and skeletal muscle CoQ_10_ measurements.

**Figure 2.**
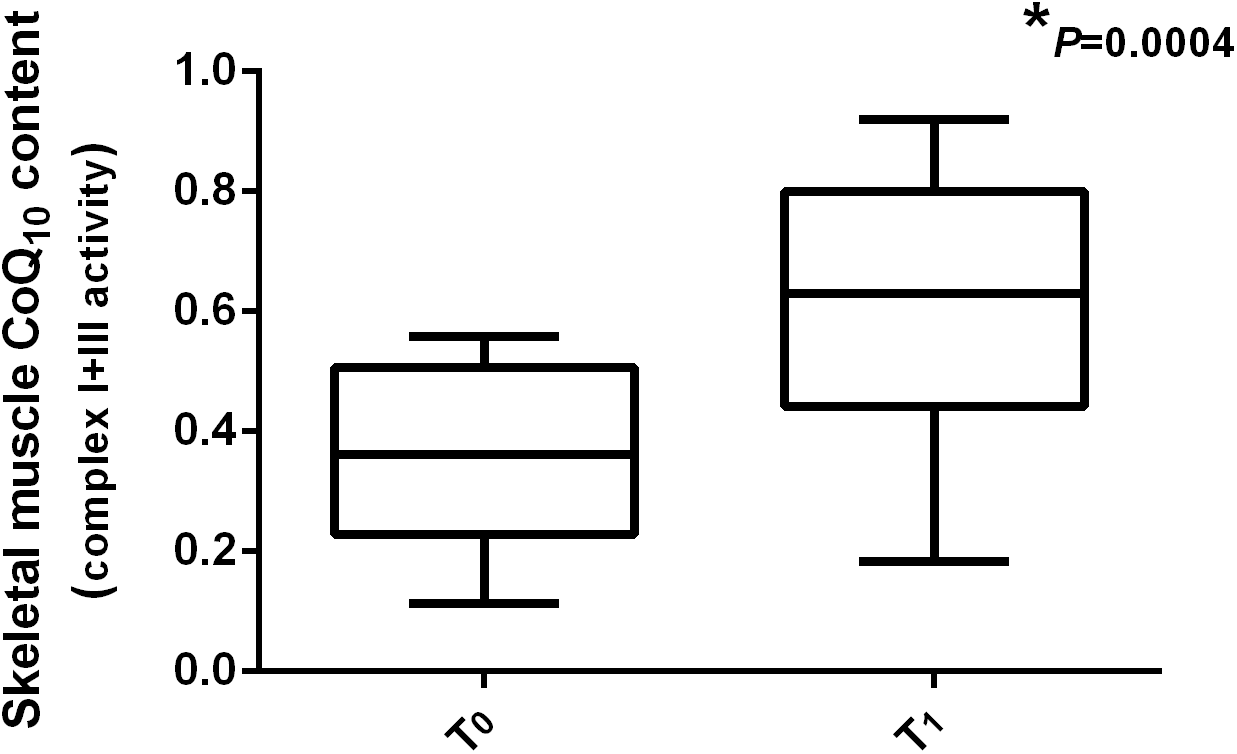
Middle gluteal skeletal muscle CoQ_10_ content for 19 young, healthy untrained Thoroughbred horses before (T_0_) and after (T_1_) 9 weeks of daily oral supplementation of the established diet with CoQ_10_ (CoQ_10_-β-cyclodextrin complex with 26% CoQ_10_, w/w). CoQ10 content was assessed by spectrophotometrically measuring skeletal muscle mitochondrial complex I+III activity. Complex I+III activity data (pmol/min/mg of muscle protein) was normalised to mitochondrial abundance (citrate synthase activity)/g of skeletal muscle. **\***Significantly different from T_0_ values (paired two-tailed Student’s *t*-test, *P*≤0.01).

## Discussion

This study demonstrated that plasma CoQ_10_ concentrations and skeletal muscle CoQ_10_ content increased in young, healthy untrained Thoroughbreds after prolonged daily oral CoQ_10_ supplementation of an established diet. In the present study, T_0_ plasma CoQ_10_ concentrations (0.13 µg/ml) were similar to a previous report evaluating 2 year-old Thoroughbreds in training (0.11 µg/ml) (Horohov et al., 2012; Sinatra, Chopra, et al., 2013; Sinatra, Jankowitz, et al., 2013), although other publications reported slightly higher basal plasma concentrations for Thoroughbreds of varying ages and fitness levels (0.19–2.1 µg/ml) (Sinatra, Chopra, et al., 2013; Topolovec et al., 2013). Following prolonged oral CoQ_10_ supplementation the mean plasma CoQ_10_ concentrations significantly increased as previously reported in studies using a similar oral cyclodextrin-CoQ_10_-based delivery system (Horohov et al., 2012; Sinatra, Chopra, et al., 2013; Sinatra, Jankowitz, et al., 2013). Intestinal absorption of CoQ_10_ has been found to be faster if CoQ_10_ is given with food (Ochiai et al., 2007) which is why we chose to supplement the horses in the morning in conjunction with their grain meals. The dose of CoQ_10_ supplementation in the present study was well tolerated by the horses with no adverse effects noted, and was chosen based on a previous study using a cyclodextrin-CoQ_10_-based delivery system (HydroQSorb, a γ-cyclodextrin [~20%] CoQ_10_ complex) (Sinatra, Chopra, et al., 2013). In humans and dogs, plasma CoQ_10_ concentrations have been found to gradually increase as the oral dosage increases (Bhagavan & Chopra, 2007), so a higher dose may have resulted in greater increases in plasma CoQ_10_ concentrations as observed in previous equine reports (Sinatra, Jankowitz, et al., 2013). However, the economic feasibility for owners and trainers were considered, as well as the fact that in humans the efficiency of oral CoQ_10_ absorption significantly decreases at extremely high doses (>300 mg) (Bhagavan & Chopra, 2007). Most researchers now believe that the formulation of oral CoQ_10_ (e.g., delivery system) is of equal if not more importance to the dosage, since this highly lipophilic molecule is typically poorly absorbed resulting in a low bioavailability despite the oral dose used as observed in humans, rats and dogs (Bank, Kagan, & Madhavi, 2011; Zhang et al., 1995; Zhang, Turunen, & Appelkvist, 1996).

CoQ_10_ is widely distributed in the body in either a reduced (i.e., ubiquinol) or oxidised (i.e., ubiquinone) form (Desbats, Lunardi, Doimo, Trevisson, & Salviati, 2015). Regardless of its form, oral CoQ_10_ is converted to ubiquinol by the enterocytes before being absorbed through the intestinal membrane, entering the systemic circulation via the lymphatic system (Bank et al., 2011) with nearly 95% of plasma CoQ_10_ present as ubiquinol (Bhagavan & Chopra, 2007). Oral CoQ_10_ bioavailability can be enhanced by altering pharmaceutical forms, with hydrophobicity of CoQ_10_ decreased by using cyclodextrin-based delivery methods (Bank et al., 2011; Jankowski et al., 2016). This delivery method significantly enhances water solubility and bioavailability (Žmitek et al., 2008) by complexing each CoQ_10_ molecule with 2 β-cyclodextrin molecules to form a water-soluble powder (Madhavi & Kagan, 2010). The oral CoQ_10_-β-cyclodextrin inclusion complex used in this study has been previously shown to be highly bioavailable with a 100% response rate in humans (e.g., all subjects had at least a doubling of plasma concentrations) and a reduced inter-subject variance (Madhavi & Kagan, 2010). High inter-subject variance is a common problem for lipophilic compounds such as CoQ_10_ because of the poor absorption, meaning that not all subjects will have the same amount of absorbance of the product (Bank et al., 2011). In the present study, there was a degree of variability between horses as has been reported for the other equine studies (Sinatra, Chopra, et al., 2013; Sinatra, Jankowitz, et al., 2013). Although budgetary restrictions meant only 12/19 horses had plasma CoQ_10_ concentrations measured, there was a 58% response rate identified when using a uniform bioavailability measurement defined as a minimum doubling of T_1_ plasma CoQ_10_ concentrations above T_0_ values.

This was a field-based study using privately-owned horses with samples not obtainable prior to the morning meal, so only the accumulation and not acute phase (0–24 hrs) of oral CoQ_10_ supplementation could be evaluated. The sample timing in the present study was based on prior studies in human subjects that demonstrated plasma concentrations to peak within 4–5 hrs of oral administration of a CoQ_10_-β-cyclodextrin inclusion complex ((Cuomo & Rabovsky, 2000; Terao et al., 2006). The CoQ_10_-β-cyclodextrin inclusion complex used in the present study has also been demonstrated in human subjects to have a sustained release resulting in the maintenance of plasma CoQ_10_ concentrations approximately 6 times higher than baseline for 24 hrs following oral administration (Madhavi & Kagan, 2010). This effect has been reported for other oral cyclodextrin complex CoQ_10_-based delivery systems, although lower sustained plasma CoQ_10_ concentrations were achieved (Terao et al., 2006).

An increase in mean skeletal muscle CoQ_10_ content was observed in the current study following prolonged oral CoQ_10_ supplementation as reflected by significant increases in CoQ_10_-dependent skeletal muscle mitochondrial function above basal activity. It has been theorised that improved CoQ_10_ absorption into the systemic circulation, elevating CoQ_10_ plasma concentrations, helps improve delivery rate into skeletal muscle (Cooke et al., 2008). The concurrent increase in plasma CoQ_10_ concentrations following supplementation found in the present study thus supports the increased skeletal muscle complex I+III activity to be a result of supplementation.

It is interesting to note that the T_1_ skeletal muscle CoQ_10_ content of three horses had fallen marginally below baseline values, potentially indicating a requirement for some horses to have higher plasma concentrations to facilitate movement of CoQ_10_ into the skeletal muscle mitochondria. There were no correlations with the degree of plasma CoQ_10_ concentrations and skeletal muscle CoQ_10_ content supporting the inability to use plasma CoQ_10_ to assess skeletal muscle CoQ_10_ content. The duration of oral CoQ_10_ supplementation has been hypothesised to contribute to the limitation of how much CoQ_10_ enters skeletal muscle mitochondria (Cooke et al., 2008). One group of researchers reported skeletal muscle CoQ_10_ content in humans to increase within 2 hours of oral supplementation, but then decrease to just above baseline values after 2 weeks of oral supplementation (Cooke et al., 2008). These researchers hypothesised that CoQ_10_ uptake into skeletal muscle may be similar to creatine monohydrate, in which there appears to be a maximal limit and/or down-regulation of transporters reached after chronic supplementation leading to a plateau and/or decrease in intramuscular content over time (Guerrero-Ontiveros & Wallimann, 1998). This warrants further investigation in the horse.

A limitation of this study was an inadequate number of horses available for a control group based on power calculations for statistical validity. A control group would have verified whether changes in dietary intake of CoQ_10_ other than supplementation contributed to the observed increases in skeletal muscle CoQ_10_ content. All study horses were housed on the same farm in adjacent pastures, with changes in grass CoQ_10_ content over the 9-week study period unlikely to have occurred since plants contain an extremely small amount of CoQ_10_, with any dietary intake for humans primarily coming from meat-based products (Kumar et al., 2009; Parmar, Jaiwal, Dhankher, & Jaiwal, 2015; Pravst, Žmitek, & Žmitek, 2010). Furthermore, even including meat-based products the typical human diet does not contain enough CoQ_10_ to significantly raise plasma CoQ_10_ concentrations above basal levels, with the daily CoQ_10_ intake from food ranging between 3–5 mg/day which is too low to significantly raise blood and tissue concentrations above basal levels (Wajda, Zirkel, & Schaffer, 2007). Since the majority of tissue CoQ_10_ content in mammals is endogenously synthesised by the mitochondria, a significantly large increase in plasma CoQ_10_ concentration is required to incite movement into tissue with human and other animal studies reporting that plasma CoQ_10_ concentrations and skeletal muscle CoQ_10_ content will not significantly increase without exogenous influences (Bhagavan & Chopra, 2007). Although plasma CoQ_10_ concentrations in humans is typically not affected by diet alone, CoQ_10_ supplementation has been shown to increase plasma CoQ_10_ concentrations, the extent of which depends upon the dosage, duration and type of formulation (Pravst et al., 2010).

It has been hypothesised that increased skeletal muscle CoQ_10_ should result in more efficient skeletal muscle energy transduction (Lenaz et al., 1999). For horses in active exercise training this may lead to improvements in responses to exercise training, delay in the onset of fatigue and enhanced recovery following intense exercise (Cooke et al., 2008; M. Kon et al., 2007; Michihiro Kon et al., 2008; Kwong et al., 2002; Mizuno et al., 2008). During exercise, movement of plasma CoQ_10_ into skeletal muscle may increase due to increased metabolic demand (M. Kon et al., 2007; Orlando et al., 2018). This theory is supported by results from a study identifying increased post-exercise intramuscular CoQ_10_ content in human athletes orally supplemented with CoQ_10_ (Cooke et al., 2008). It has recently been shown that resting skeletal muscle CoQ_10_ content is associated with *myostatin* (*MSTN*) genotype (SNP g.66493737C>T) in untrained Thoroughbred horses (Rooney, Porter, Katz, & Hill, 2017). ETC combined complex I+III and II+III activities (indirect measures of CoQ_10_ content) were significantly lower in resting skeletal muscle from TT *MSTN* genotype horses as compared to CT and CC horses. In this same study, restoration of complex I+III and II+III activity was achieved following *in vitro* supplementation with exogenous coenzyme Q_1_. Based on the observed differences in basal concentrations of skeletal muscle CoQ_10_ between *MSTN* genotypes in Thoroughbreds, oral supplementation with CoQ_10_ may have a greater efficacy in skeletal muscle of horses with the TT *MSTN* genotype, especially for TT horses training and competing in endurance-related competitions. In the present study, the number of horses with different *MSTN* genotypes was too small to assess for genotype-specific variation in plasma and skeletal muscle CoQ_10_ concentrations after supplementation, but this certainly warrants further investigation.

### Conclusion

In summary, this study demonstrates that prolonged daily oral supplementation of a grass and oat diet of young, healthy untrained Thoroughbreds with a CoQ_10_-β-cyclodextrin inclusion complex significantly increases mean plasma concentration and skeletal muscle CoQ_10_ content, although a degree of variability was identified for some horses. Additional research is warranted to investigate the effects of *MSTN* genotype, training and exercise on skeletal muscle CoQ_10_ content in CoQ_10_-β-cyclodextrin inclusion complex supplemented and un-supplemented Thoroughbreds.

## Acknowledgements

The authors wish to thank the owners of the horses used in the study and the farm staff for the assistance provided during the study. The authors would also like to acknowledge the assistance provided by Plusvital Ltd. staff. This study has emanated from research conducted with the financial support from an Enterprise Ireland Innovation Partnership Programme Grant (IP/2016/0503).

## References

Alf, D., Schmidt, M. E., & Siebrecht, S. C. (2013). Ubiquinol supplementation enhances peak power production in trained athletes: a double-blind, placebo controlled study. Journal of the International Society of Sports Nutrition, 10(1), 24.

Bank, G., Kagan, D., & Madhavi, D. (2011). Coenzyme Q10: clinical update and bioavailability. Journal of Evidence-Based Complementary & Alternative Medicine, 16(2), 129–137.

Bhagavan, H. N., & Chopra, R. K. (2007). Plasma coenzyme Q10 response to oral ingestion of coenzyme Q10 formulations. Mitochondrion, 7, S78–S88.

Bonetti, A., Solito, F., Carmosino, G., Bargossi, A., & Fiorella, P. (2000). Effect of ubidecarenone oral treatment on aerobic power in middle-aged trained subjects. Journal of Sports Medicine and Physical Fitness, 40(1), 51.

Carroll, C., & Huntington, P. (1988). Body condition scoring and weight estimation of horses. Equine Veterinary Journal, 20(1), 41–45.

Cooke, M., Iosia, M., Buford, T., Shelmadine, B., Hudson, G., Kerksick, C., … Willoughby, D. (2008). Effects of acute and 14-day coenzyme Q10 supplementation on exercise performance in both trained and untrained individuals. Journal of the International Society of Sports Nutrition, 5(1), 1.

Cuomo, J., & Rabovsky, A. (2000). Comparative Bioavailability of Coenzyme Q10 in Four Formulations. USANA Clinical Research Bulletin Number, 5.

Desbats, M. A., Lunardi, G., Doimo, M., Trevisson, E., & Salviati, L. (2015). Genetic bases and clinical manifestations of coenzyme Q10 (CoQ10) deficiency. Journal of Inherited Metabolic Disease, 38(1), 145–156.

Duncan, A. J., Heales, S. J., Mills, K., Eaton, S., Land, J. M., & Hargreaves, I. P. (2005). Determination of coenzyme Q10 status in blood mononuclear cells, skeletal muscle, and plasma by HPLC with di-propoxy-coenzyme Q10 as an internal standard. Clinical Chemistry, 51(12), 2380–2382.

Greenberg, S., & Frishman, W. H. (1990). Co-enzyme Q10: a new drug for cardiovascular disease. The Journal of Clinical Pharmacology, 30(7), 596–608.

Guerrero-Ontiveros, M. L., & Wallimann, T. (1998). Creatine supplementation in health and disease. Effects of chronic creatine ingestion in vivo: down-regulation of the expression of creatine transporter isoforms in skeletal muscle. Molecular and Cellular Biochemistry, 184(1-2), 427–437.

Hofman-Bang, C., Rehnqvist, N., Swedberg, K., Wiklund, I., & Åström, H. (1995). Coenzyme Q10 as an adjunctive in the treatment of chronic congestive heart failure. Journal of Cardiac Failure, 1(2), 101–107.

Horohov, D. W., Sinatra, S. T., Chopra, R. K., Jankowitz, S., Betancourt, A., & Bloomer, R. J. (2012). The effect of exercise and nutritional supplementation on proinflammatory cytokine expression in young racehorses during training. Journal of Equine Veterinary Science, 32(12), 805–815.

Jankowski, J., Korzeniowska, K., Cieślewicz, A., & Jabłecka, A. (2016). Coenzyme Q10–A new player in the treatment of heart failure? Pharmacological Reports, 68(5), 1015–1019.

Kaikkonen, J., Tuomainen, T.-P., Nyyssönen, K., & Salonen, J. T. (2002). Coenzyme Q10: absorption, antioxidative properties, determinants, and plasma levels. Free Radical Research, 36(4), 389–397.

Kamzalov, S., Sumien, N., Forster, M. J., & Sohal, R. S. (2003). Coenzyme Q intake elevates the mitochondrial and tissue levels of coenzyme Q and α-tocopherol in young mice. Journal of Nutrition, 133(10), 3175–3180.

Kon, M., Kimura, F., Akimoto, T., Tanabe, K., Murase, Y., Ikemune, S., & Kono, I. (2007). Effect of Coenzyme Q10 supplementation on exercise-induced muscular injury of rats. Exercise Immunology Review, 13, 76–88.

Kon, M., Tanabe, K., Akimoto, T., Kimura, F., Tanimura, Y., Shimizu, K., … Kono, I. (2008). Reducing exercise-induced muscular injury in kendo athletes with supplementation of coenzyme Q 10. British Journal of Nutrition, 100(04), 903–909.

Kumar, A., Kaur, H., Devi, P., & Mohan, V. (2009). Role of coenzyme Q10 (CoQ10) in cardiac disease, hypertension and Meniere-like syndrome. Pharmacology & Therapeutics, 124(3), 259–268.

Kwong, L. K., Kamzalov, S., Rebrin, I., Bayne, A.-C. V., Jana, C. K., Morris, P., … Sohal, R. S. (2002). Effects of coenzyme Q10 administration on its tissue concentrations, mitochondrial oxidant generation, and oxidative stress in the rat. Free Radical Biology and Medicine, 33(5), 627–638.

Ledwith, A., & McGowan, C. M. (2004). Muscle biopsy: a routine diagnostic procedure. Equine Veterinary Education, 16(2), 62–67.

Leelarungrayub, D., Sawattikanon, N., Klaphajone, J., Pothongsunan, P., & Bloomer, R. J. (2010). Coenzyme Q10 supplementation decreases oxidative stress and improves physical performance in young swimmers: a pilot study. The Open Sports Medicine Journal, 4(1), 1–8.

Lenaz, G., Fato, R., Di Bernardo, S., Jarreta, D., Costa, A., Genova, M. L., & Castelli, G. P. (1999). Localization and mobility of coenzyme Q in lipid bilayers and membranes. Biofactors, 9(2-4), 87–93.

Lerman-Sagie, T., Rustin, P., Lev, D., Yanoov, M., Leshinsky-Silver, E., Sagie, A., … Munnich, A. (2001). Dramatic improvement in mitochondrial cardiomyopathy following treatment with idebenone. Journal of inherited metabolic disease, 24(1), 28–34.

Leshinsky-Silver, E., Levine, A., Nissenkorn, A., Barash, V., Perach, M., Buzhaker, E., … Lerman-Sagie, T. (2003). Neonatal liver failure and Leigh syndrome possibly due to CoQ-responsive OXPHOS deficiency. Molecular genetics and metabolism, 79(4), 288–293.

Linnane, A. W., Kopsidas, G., Zhang, C., Yarovaya, N., Kovalenko, S., Papakostopoulos, P., … Richardson, M. (2002). Cellular redox activity of coenzyme Q10: effect of CoQ 10 supplementation on human skeletal muscle. Free Radical Research, 36(4), 445–453.

Madhavi, D., & Kagan, D. (2010). A study on the bioavailability of a novel sustained-release coenzyme Q10-β-cyclodextrin complex. Integrative Medicine, 9(1), 20–24.

Mas, E., & Mori, T. A. (2010). Coenzyme Q10 and Statin Myalgia: What is the Evidence?. Curr Atheroscler Rep, 12(6), 407–413.

Mizuno, K., Tanaka, M., Nozaki, S., Mizuma, H., Ataka, S., Tahara, T., … Kuratsune, H. (2008). Antifatigue effects of coenzyme Q10 during physical fatigue. Nutrition, 24(4), 293–299.

Mortensen, S. A., Rosenfeldt, F., Kumar, A., Dolliner, P., Filipiak, K. J., Pella, D., … investigators, Q.-S. s. (2014). The effect of coenzyme Q10 on morbidity and mortality in chronic heart failure: results from Q-SYMBIO: a randomized double-blind trial. J Am Coll Cardiol HF, 2(6), 641–649.

Ochiai, A., Itagaki, S., Kurokawa, T., Kobayashi, M., Hirano, T., & Iseki, K. (2007). Improvement in intestinal coenzyme Q10 absorption by food intake. Yakugaku Zasshi. Journal of the Pharmaceutical Society of Japan, 127(8), 1251–1254.

Olson, R. E., & Rudney, H. (1983). Biosynthesis of ubiquinone. Vitamins & Hormones, 40, 1–43.

Orlando, P., Silvestri, S., Galeazzi, R., Antonicelli, R., Marcheggiani, F., Cirilli, I., … Tiano, L. (2018). Effect of ubiquinol supplementation on biochemical and oxidative stress indexes after intense exercise in young athletes. Redox Report, 23(1), 136–145.

Parmar, S. S., Jaiwal, A., Dhankher, O. P., & Jaiwal, P. K. (2015). Coenzyme Q10 production in plants: current status and future prospects. Critical Reviews in Biotechnology, 35(2), 152–164.

Powers, W. J., Haas, R. H., Le, T., Videen, T. O., Hershey, T., McGee-Minnich, L., & Perlmutter, J. S. (2007). Normal platelet mitochondrial complex I activity in Huntington’s disease. Neurobiology of Disease, 27(1), 99–101.

Pravst, I., Žmitek, K., & Žmitek, J. (2010). Coenzyme Q10 contents in foods and fortification strategies. Critical Reviews in Food Science and Nutrition, 50(4), 269–280.

Rooney, M. F., Porter, R. K., Katz, L. M., & Hill, E. W. (2017). Skeletal muscle mitochondrial bioenergetics and associations with myostatin genotypes in the Thoroughbred horse. PloS one, 12(11), e0186247. doi: 10.1371/journal.pone.0186247

Sinatra, S. T., Chopra, R. K., Jankowitz, S., Horohov, D. W., & Bhagavan, H. N. (2013). Coenzyme Q10 in equine serum: response to supplementation. Journal of Equine Veterinary Science, 33(2), 71–73.

Sinatra, S. T., Jankowitz, S. N., Chopra, R. K., & Bhagavan, H. N. (2013). Plasma coenzyme Q10 and tocopherols in thoroughbred race horses: Effect of coenzyme Q10 supplementation and exercise. Journal of Equine Veterinary Science, 34(2), 265–269.

Smith, P. K., Krohn, R. I., Hermanson, G. T., Mallia, A. K., Gartner, F. H., Provenzano, M. D., … Klenk, D. C. (1985). Measurement of protein using bicinchoninic acid. Analytical Biochemistry, 150(1), 76–85.

Srere, P. A. (1969). [1] Citrate synthase: [EC 4.1.3.7. Citrate oxaloacetate-lyase (CoA-acetylating)]. In M. L. John (Ed.),. Methods in Enzymology (Vol. Volume 13, pp. 3–11): Academic Press.

Svensson, M., Malm, C., Tonkonogi, M., Ekblom, B., Sjödin, B., & Sahlin, K. (1999). Effect of Q10 supplementation on tissue Q10 levels and adenine nucleotide catabolism during high-intensity exercise. International Journal of Sport Nutrition, 9(2), 166–180.

Terao, K., Nakata, D., Fukumi, H., Schmid, G., Arima, H., Hirayama, F., & Uekama, K. (2006). Enhancement of oral bioavailability of coenzyme Q10 by complexation with γ-cyclodextrin in healthy adults. Nutrition Research, 26(10), 503–508.

Topolovec, M. B., Kruljc, P., Prošek, M., Križman, P. J., Šmidovnik, A., & Svete, A. N. (2013). Endogenous plasma coenzyme Q10 concentration does not correlate with plasma total antioxidant capacity level in healthy untrained horses. Research in Veterinary Science, 95(2), 675–677.

Tran, U. C., & Clarke, C. F. (2007). Endogenous synthesis of coenzyme Q in eukaryotes. Mitochondrion, 7, S62–S71.

Turunen, M., Swiezewska, E., Chojnacki, T., Sindelar, P., & Dallner, G. (2002). Regulatory aspects of coenzyme Q metabolism. Free Radical Research, 36(4), 437–443.

Wajda, R., Zirkel, J., & Schaffer, T. (2007). Increase of bioavailability of coenzyme Q10 and vitamin E. Journal of Medicinal Food, 10(4), 731–734.

Yang, Y.-K., Wang, L.-P., Chen, L., Yao, X.-P., Yang, K.-Q., Gao, L.-G., & Zhou, X.-L. (2015). Coenzyme Q10 treatment of cardiovascular disorders of ageing including heart failure, hypertension and endothelial dysfunction. Clinica Chimica Acta, 450, 83–89.

Zhang, Y., Aberg, F., & Appelkvist, E.-L. (1995). Uptake of Dietary Coenzyme Q Supplement ls limited in rats. The Journal of Nutrition, 125(3), 446–453.

Zhang, Y., Turunen, M., & Appelkvist, E.-L. (1996). Restricted uptake of dietary coenzyme Q is in contrast to the unrestricted uptake of alpha-tocopherol into rat organs and cells. The Journal of Nutrition, 126(9), 2089.

Zhou, S., Zhang, Y., Davie, A., & Marshall-Gradisnik, S. (2005). Muscle and plasma coenzyme Q10 concentration, aerobic power and exercise economy of healthy men in response to four weeks of supplementation. Journal of Sports Medicine and Physical Fitness, 45(3), 337.

Žmitek, J., Šmidovnik, A., Fir, M., Prošek, M., Žmitek, K., Walczak, J., & Pravst, I. (2008). Relative bioavailability of two forms of a novel water-soluble coenzyme Q10. Annals of Nutrition and Metabolism, 52(4), 281–287.

